# A simple self-decoding model for neural coding

**DOI:** 10.1101/2022.02.12.480019

**Authors:** Thach V. Bui

## Abstract

Neural coding is an important tool to discover the inner workings of mind. In this work, we propose and consider a simple but novel self-decoding model for neural coding based on the principle that the neuron body represents ongoing stimulus while dendrites are used to store that stimulus as a memory. In particular, suppose *t* spiking presynaptic neurons transmit any stimulus directly to a population of *n* postsynaptic neurons, a postsynaptic neuron spikes if it does not connect to an inhibitory presynaptic neuron, and every stimulus is represented by up to *d* spiking postsynaptic neurons.

Our hypothesis is that the brain is organized to functionally satisfy the following six criteria: (i) decoding objective, i.e., there are up to *r*−1 ≥ 0 additional spiking postsynaptic neurons in response to a stimulus along with the spiking postsynaptic neurons representing the stimulus, (ii) smoothness, i.e., similar stimuli are encoded similarly by the presynaptic neurons, (iii) optimal information transmission, i.e., *t* is minimized, (iv) optimal energetic cost, i.e., only the *t* presynaptic neurons and the postsynaptic neurons representing a stimulus spike, (v) low-dimensional representation, i.e., *d* = *o*(*n*), and (vi) sparse coding, i.e., *t* = *o*(*n*).

Our finding is that some criteria cause or correlate with others. Let the characteristic set of a postsynaptic neuron be the set of the presynaptic neurons it connects with. We prove that (i) holds *if and only if* the union of the *r* characteristic sets of any *r* postsynaptic neurons is not included in the union of the *d* characteristic sets of *d* other postsynaptic neurons. Consequently, (ii) is attained. More importantly, we suggest that the decoding objective (i) and optimal information transmission (iii) play a fundamental role in neural computation, while (v) and (vi) correlate to each other and correlate with (iii) and (iv). We examine our hypothesis by statistically testing functional connectivity network and the presynaptic-postsynaptic connectivity in layer 2 of the medial entorhinal cortex of a rat.

## I. Introduction

Since the discovery that a neuron either spikes or does not spike in response to a stimulus [1], neural coding, i.e., the neural representation of information, could have been considered as discrete events given a population of neurons. It then has been extensively studied to have a better understanding on how the external world is reflected and processed in the brain.

Let *t* and *n* be the number of presynaptic neurons and the number of postsynaptic neurons, respectively. When studying neurons in response to stimuli, the following six criteria are always or usually observed: (i) decoding objective, i.e., there are up to *r*−1 ≥ 0 additional spiking postsynaptic neurons in response to the stimulus along with the spiking postsynaptic neurons representing the stimulus, (ii) smoothness [2], [3], i.e., similar stimuli encoded similarly by the presynaptic neurons, (iii) optimal information transmission [4]–[6], i.e., *t* is minimized, (iv) optimal energetic cost [7]–[9], i.e., only the *t* presynaptic neurons and the postsynaptic neurons representing a stimulus spike, (v) low-dimensional representation [10]–[15], i.e., *d* = *o*(*n*), and (vi) sparse coding [16]–[19], i.e., *t* = *o*(*n*). Note that the first criteria should be held in brain. Otherwise, one cannot distinguish different stimuli. Exact decoding occurs when *r* = 1, whereas approximate decoding occurs when *r* ≥ 2. Note that approximate decoding could imply exact decoding if some special conditions hold.

There are a vast number of works that try to explain the above six criteria. There are four representative models that are considered to match reality, such as time-dependent firing rate [20], temporal coding [21]–[25], population coding [26]–[28], and sparse coding, which is suggested to be a general strategy of neural systems to increase memory capacity, [16]–[19]. Since information theory is a useful tool to maximize information transmission and minimize the redundancy of stimulus representations, it has been received substantial attention and influence on theoretical neuroscience [29]–[32]. In contrast, coding theory, a research field whose aim is to engineer codes in practice, has been considered less for neural coding. However, recent studies [33]–[35] show that combinatorial neural codes, which is a branch of coding theory, have advantages in representing stimuli. Recently, Chaudhuri and Fiete [36] propose a model based on expander codes to attain the two capabilities of exponential capacity and high robustness to noise with a self-decoding algorithm. They then use that model to generate artificial neural networks to test their hypotheses. Followed that work, Maoz et al. [37] proposed a neural code based on the sparse and random connectivity of real neural circuits to estimate spiking patterns in brain. Both works suggest that random sparse connectivity, i.e., criterion (vi), is a key design principle for neural computation. However, in our work, we suggest an alternative conclusion. To our knowledge, no existing neural network model can be proved to be the right morphology of brain, satisfies all six criteria, or has been used to investigate relationships between the six criteria mathematically.

### Contributions

In this work, we propose a model that is consistent with the principle that the neuron body represents ongoing stimulus while dendrites are used to store that stimulus as a memory. Consider the following model. Suppose that every stimulus is represented by up to *d* spiking postsynaptic neurons, every presynaptic neuron spikes, the postsynaptic neurons are activated directly by the presynaptic neurons, and a postsynaptic neuron spikes if it does not connect to an inhibitory presynaptic neuron. We make a hypothesis that brain is organized to functionally satisfy the above six criteria. Our finding is that some criteria cause or correlate others. In particular, we suggest that the decoding objective (i) and optimal information transmission (iii) play a fundamental role in neural computation while (v) and (vi) correlate to each other and correlate with (iii) and (iv). Finally, we validate our hypothesis by statistically testing real datasets.

#### Theoretical results

Let the characteristic set of a postsynaptic neuron be the set of the presynaptic neurons it connects with. We prove that the decoding objective holds *if and only if* the union of the *r* characteristic sets of any *r* postsynaptic neurons is not included in the union of the *d* characteristic sets of *d* other postsynaptic neurons. To our knowledge, our work is the first to prove that the brain morphology must have a certain topology under certain conditions. Smoothness and the optimal energetic cost are direct consequences of that morphology. We also show that exact coding and optimal information transmission cause the optimal energetic cost to encode and decode a stimulus. Moreover, when *r* = *d*, the required number of presynaptic neurons to encode and decode a stimulus meets the information-theoretic bound by allowing up to *d*−1 additional spiking postsynaptic neurons along with the postsynaptic neurons representing that stimulus. We then show that the low-dimensional representation has correlations with optimal information transmission and sparse coding. These findings are summarized in Figure 1.

**Fig. 1.**
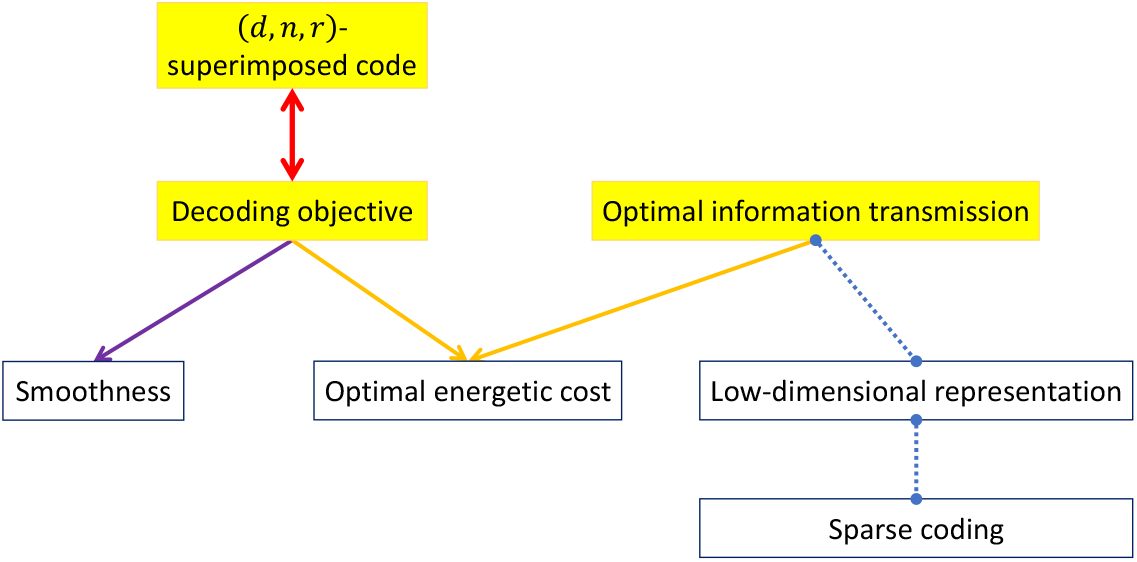
Relationships between six criteria in brain. The sources of incoming arrows of a box cause the criterion in that box, while the dashed line with two round points represents the correlation between the two ends. The decoding objective is equivalent to a superimposed (*d, n, r*)-code (defined later). Moreover, the decoding objective causes smoothness. Exact decoding of the decoding objective and optimal information transmission cause the optimal energetic cost. If the decoding objective and optimal information transmission are satisfied, the low-dimensional representation causes sparse coding when 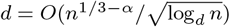 with *α <* 1*/*3, and sparse coding also causes the low-dimensional representation. Moreover, the low-dimensional representation correlates with optimal information transmission.

#### Experimental results

Let *M* = (*m*_*ij*_) be a presynaptic-postsynaptic connectivity matrix, where *m*_*ij*_ = 1 means the presynaptic neuron *i* has a synaptic connection with the postsynaptic neuron *j*, and *m*_*ij*_ = 0 otherwise. Theoretically, with the proposed model, to obtain the decoding objective, *M* must be a superimposed code (defined later), and vice versa. However, *M might not be* the superimposed code due to the following reasons: the proposed model may not be fit in practice, all presynaptic and postsynaptic neurons are not observed, and some errors occur when constructing *M*. Therefore, to verify whether *M* is a superimposed code, one can the probability of *M* being a superimposed code, i.e., the probability of matrix matching, by using a population proportion. From the relationships described in Figure 1 and our analysis in Section III-C, our hypothesis is true if the proposed model is satisfied, *M* is a superimposed code, the low-dimensional representation and optimal information transmission hold. Therefore, if the proposed model is satisfied, by varying *d* and setting *t* as optimal information transmission accordingly such that *t* = *o*(*n*), the probability of matrix matching, is equal to the probability of hypothesis matching. Note that there are two types of superimposed codes, namely disjunct matrices and selectors (defined later), that are equivalent to exact decoding and approximate decoding, respectively.

We examine our hypothesis with two real datasets, which are a functional connectivity network and the presynaptic-postsynaptic connectivity in layer 2 of the medial entorhinal cortex of a rat. In the first dataset, we can assume that the proposed model is satisfied. Hence, the probability of hypothesis matching is the probability of matrix matching. When *d* is small and *t/n* ≤ 0.276, the probability of hypothesis matching converges quickly to 1. Once *d* grows, the probability that *M* is a disjunct matrix (exact decoding) still converges to 1, however, not as fast as the probability that *M* is a selector (approximate decoding). For the second dataset, *M* has size of 18 × 180. We only test the probability of matrix matching for matrix *M*. Although this dataset is small, *M* still is almost a disjunct matrix or a selector for *d* = 2.

The above results suggests two possibilities: 1) for most sets of postsynaptic neurons, they have the structure of disjunct matrices to allow exact decoding; and 2) approximate decoding is used first to estimate stimuli before processing exact decoding. Moreover, it highly suggests that *selectors (defined later) may be the right topology of the presynaptic-postsynaptic connectivity in brain*.

## II. Proposed Model

In this section, we first provide an overview of biological neural networks and their properties, and then mathematically model them.

### A. Biological neural networks

#### 1) Settings

Biological neural networks are present in a way that can be used to process stimuli efficiently. Here we make three basic assumptions on presynaptic neurons, postsynaptic neurons, and the presynaptic-postsynaptic connectivity to serve that purpose.

### Combinatorial representation setting

Given a population of *n* postsynaptic neurons, suppose that every stimulus is represented by up to *d* spiking postsynaptic neurons.

### Decoding rule

We make two assumptions on spiking for presynaptic and postsynaptic neurons, which are also naturally a self-decoding procedure in brain, as follows:

1. Every presynaptic neuron spikes.
2. A postsynaptic neuron spikes if it does not connect to an inhibitory presynaptic neuron.

For simplicity, we set every presynaptic neuron as a hybrid neuron [38], [39], i.e., it can behave either as an excitatory neuron or an inhibitory neuron.

### 1-stage design

To induce dendritic spikes, near-synchronous activation [22] or tight synchronicity [23] of spiking presynaptic neurons are often observed in the apical tuft or apical trunk and apical oblique dendrites. Therefore, the latency to propagate spikes from the presynaptic neurons to the postsynaptic neurons should be taken into account. The optimal latency happens when the postsynaptic neurons are activated directly by the presynaptic neurons. This is equivalent to a one-layer feedforward network [40]. Thus, we assume that the optimal latency is obtained here.

#### 2) Representation

We now formulate stimulus and its information generated at the presynaptic neurons. Given a set of *n* postsynaptic neurons labeled {1, 2, …, *n*} = [*n*], let 𝒳 ⊆ {0, 1} ^*n*^ be the discretized stimulus space (the ambient space) representing for all stimuli. For any vector **v** = (*v*_1_, …, *v*_*n*_) ∈ {0, 1}^*n*^, *v*_*j*_ = 1 means the postsynaptic neuron *j* spikes and *v*_*j*_ = 0 means otherwise. Let **x** = (*x*_1_, …, *x*_*n*_) ∈ 𝒳 be the binary representation vector for an input stimulus.

Given a *fixed* set of *t* presynaptic neurons labeled {1, 2, …, *t}* = [*t*], let 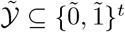 be the encoded space of the ambient space. For a vector 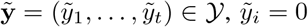 means the presynaptic neuron *i* is inhibitory and spiking, and 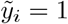 means the presynaptic neuron *i* is excitatory and spiking.

A stimulus represented by vector **x** is encoded at the *t* presynaptic neurons as a vector 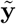. These presynaptic neurons spike and activate the postsynaptic neurons that represent the stimulus. We illustrate this stimulus encoding and decoding procedure in Figure 2. The input stimulus is represented by *n* = 12 postsynaptic neurons with the neurons 3 and 9 spiking. This stimulus is encoded by *t* = 9 presynaptic neurons in which the presynaptic neurons 1, 3, 4, 7, and 8 behaves as excitatory neurons while the presynaptic 2, 5, 6, and 9 behaves as inhibitory neurons. Follow the spiking rules of the presynaptic neurons, the two postsynaptic neurons 3 and 9 spike and therefore the stimulus is written at the postsynaptic neurons as intended.

**Fig. 2.**
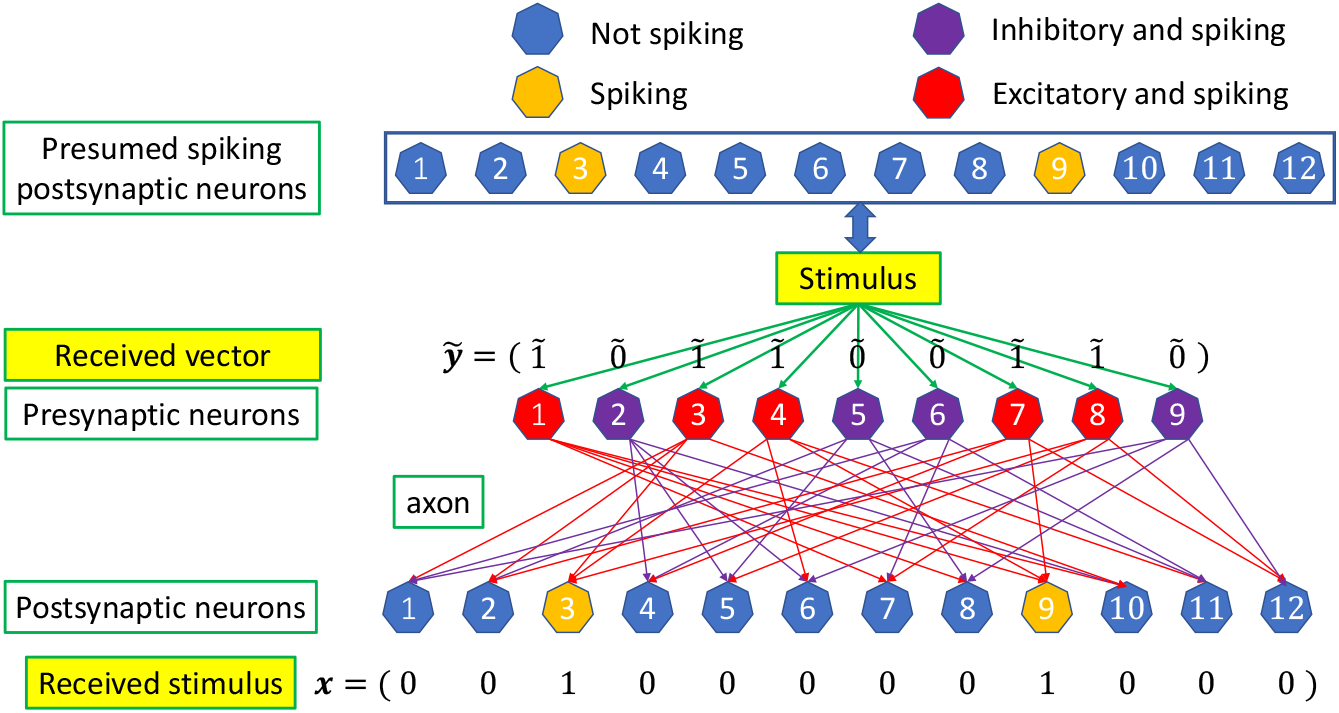
Stimulus encoding and decoding procedure in biological neural networks. An input stimulus is represented by 12 postsynaptic neurons with the neurons 3 and 9 spiking. A postsynaptic neuron filled with orange (respectively, blue) is (respectively, not) spiking. The stimulus is encoded by 9 presynaptic neurons in which the presynaptic neurons 1, 3, 4, 7, and 8 behaves as excitatory neurons while the presynaptic 2, 5, 6, and 9 behaves as inhibitory neurons. A presynaptic neuron filled with red (respectively, purple) is excitatory (respectively, inhibitory) and spiking. The postsynaptic neurons are spiked by the presynaptic neurons via the presynaptic axons, which are represented by line arrows.

Sheffield and Dombeck [41] suggest that the neuron body represents ongoing stimulus while dendrites are used to store that stimulus as a memory. The second argument is widely supported by several studies [42]–[46]. Our task now is to describe the stimulus encoding and decoding procedure here in a mathematical way such that it is consistent with the way neuroscientists usually think about neural coding. The details are presented in the next subsection.

### B. General model

We present the general idea of the proposed model here and illustrate it in Figure 3. First, we discretize the stimulus space and the encoding map space.

**Fig. 3.**
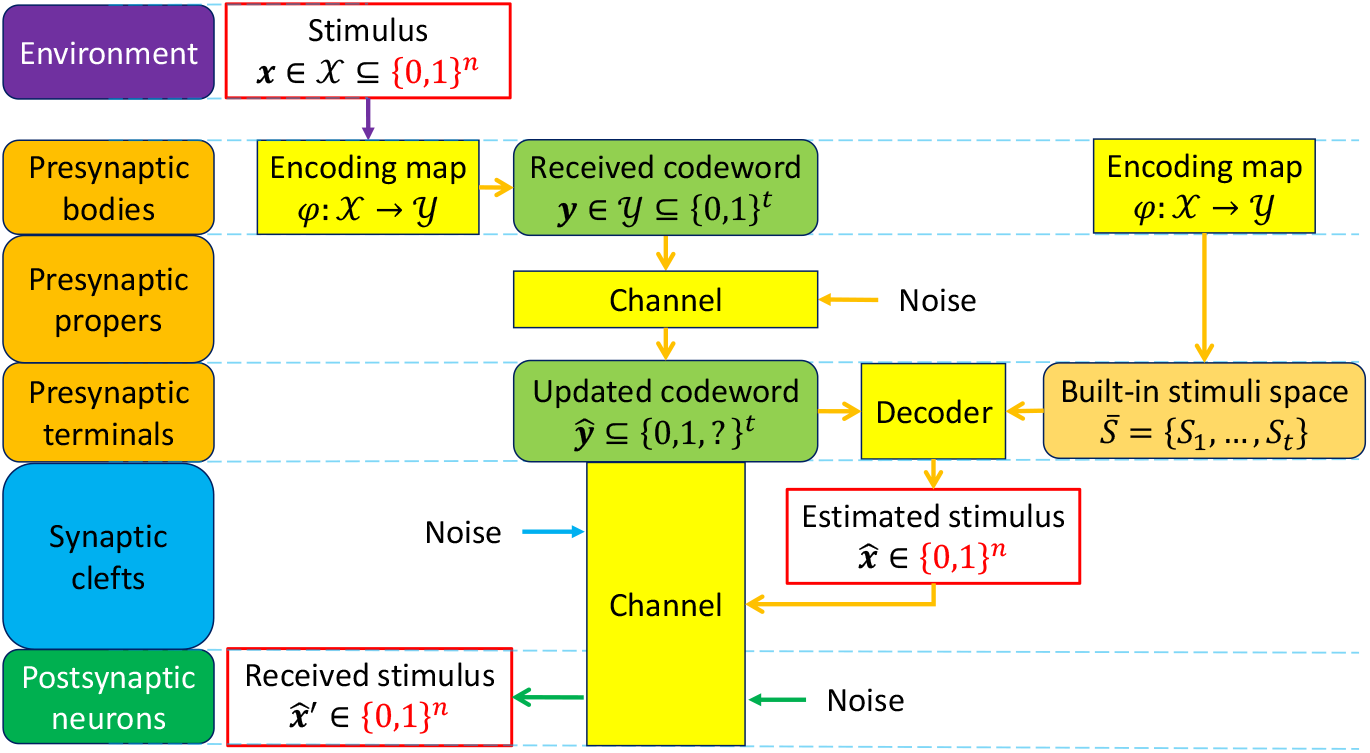
General model for stimulus encoding and decoding. A stimulus **x** ∈ 𝒳 is encoded by a codeword **y** = *φ*(**x**) ∈ 𝒴 via the encoding map *φ*. The received codeword **y** ∈ 𝒴 is then transmitted through the (noisy) channel located at the presynaptic propers to reach the presynaptic terminals and becomes an updated codeword **ŷ** = (*ŷ*_1_, …, *ŷ*_*t*_) ∈ {0, 1, ?}^*t*^. The decoder takes the updated codeword **ŷ** and the built-in stimulus space 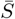 as two inputs to produce an estimated stimulus 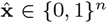 of the stimulus **x**. The estimated stimulus then goes through a (noisy) channel located at the postsynaptic neurons and the synaptic clefts to finally generate the received stimulus 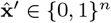 at the presynaptic neurons.

Given a *fixed* set of *t* presynaptic neurons labeled {1, 2, …, *t}* = [*t*], let 𝒴 ⊆ {0, 1}^*t*^ be the encoded space of the ambient space and be called *the code*. The elements of the code are called *codewords*.

Suppose there is a function *φ* that maps 𝒳 to 𝒴, i.e., *φ* : 𝒳 → 𝒴. In particular, for each stimulus **x** ∈ 𝒳, there exists a codeword **y** ∈ 𝒴 such that **y** = *φ*(**x**). The encoding map locates at the presynaptic bodies.

Let 𝒮_*i*_ be the set of all postsynaptic neurons that the presynaptic *i*th neuron innervates. Then we define 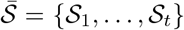 as a built-in stimulus space which is independent to most of ongoing stimuli but is subjected to be gradually changed over time due to the ongoing stimuli (plasticity). This space is built based on the encoding map *φ*.

We describe the full process of encoding and decoding for a stimulus and locations that each part of our model occurs at here. A stimulus **x** ∈ 𝒳 is represented by a codeword **y** = *φ*(**x**) ∈ 𝒴 via the encoding map *φ*. The encoding map and the received codeword **y** are located at the presynaptic bodies. The received codeword **y** is then transmitted through the channel located at the presynaptic axon (presynaptic propers) to reach the presynaptic terminals. Under the effect of noise in the channel, it becomes an updated codeword **ŷ** = (*ŷ* _1_, …, *ŷ* _*t*_) ∈ { 0, 1, ?}^*t*^. The decoder takes the updated codeword **ŷ** and the built-in stimulus space 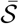 as two inputs to produce an estimated stimulus 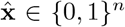 of the stimulus **x** with the following decoding rules:

1. Every postsynaptic neuron spikes, except for those belonging to 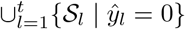.
2. If *ŷ*_*i*_ =? then every postsynaptic neuron in 𝒮_*i*_ is not affected by the presynaptic neuron *i*.

The first rule can be interpreted as follows: if *ŷ*_*i*_ = 0 then every postsynaptic neuron in 𝒮_*i*_ does not spike and if *ŷ*_*i*_ = 1 then every postsynaptic neuron in 𝒮_*i*_ spikes, except for those belonging to 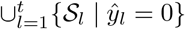.

The updated codeword, the decoder, and the built-in stimulus space are located at the presynaptic terminals while the estimated stimulus is located at the synaptic clefts. The estimated stimulus then goes through a (noisy) channel located at the postsynaptic neurons and the synaptic clefts to finally register the received stimulus 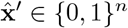 at the presynaptic neurons.

Our proposed model is consistent with the representation in Section II-A2 and the two arguments of Sheffield and Dombeck [41]. In our model, the output of the encoding of a stimulus into a pattern of neural activity is the received codeword, as illustrated in the flow from the stimulus **x** to the received codeword **y**. This is consistent with the first argument that the neuron body represents ongoing stimulus. Moreover, the postsynaptic dendrites are only used to store the stimulus as a memory in our model because the output of the decoder is located at the synaptic clefts. This then coincides with the second argument.

### C. Simplified model in the noiseless setting

As suggested by Sheffield and Dombeck [41], the postsynaptic neurons are just a place to store stimuli as memory. This means the procedure to get the received stimulus from the estimated stimulus does not have any impact on the stimulus encoding and decoding procedure. Furthermore, Schneidman et al. [34] suggest that combinatorial coding in terms of synergy might carry redundant information which is used to attain noise-resilience. We therefore can consider the stimulus encoding and decoding procedure with the pathway from the stimulus in environment to the estimated stimulus without noise. This model is called the *simplified model in the noiseless setting* which is *our focus* throughout this work. In particular, an input stimulus **x** ∈ 𝒳 ⊆ {0, 1}^*n*^ with |`supp`(**x**)| ≤ *d* goes through the encoding map *φ* : 𝒳 → 𝒴 ⊆ {0, 1}^*t*^ to output a received codeword **y** = *φ*(**x**) ∈ 𝒴, which is the same as the updated codeword, where `supp`(**x**) = {*j* ∈ [*n*] | *x*_*j*_ = 1}. The received codeword and the built-in stimulus space then go through the decoder to produce an estimated vector of the input stimulus 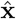, which is ideally the same as the input stimulus **x**. Note that the decoding rule is every postsynaptic neuron spikes, except for those belonging to 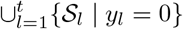.

## III. Theoretical results

In this section, we formally define the six criteria mentioned in Section I. We show that to decode a stimulus with a few or with no errors, the brain must have a certain morphology. Then we analyze relationships between the six criteria in terms of causality and correlation.

### A. Criteria formulation

Let `supp`(**v**) = {*j* ∈ [*p*] | *v*_*j*_ = 1} for **v** = (*v*_1_, …, *v*_*p*_). We first define the decoding objective for a stimulus.

#### Criterion 1 (Decoding objective).

*We require that there are up to r* − 1 *additional postsynaptic neurons along with the postsynaptic neurons representing a stimulus spiking in response to that stimulus. In other words*, 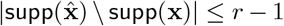 *and* 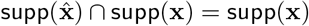. *When r >* 1, *the decoding is approximate, whereas the decoding is exact when r* = 1.

Smoothness is an important property of brain to represent a large number of stimuli [2]. A neural code is smooth if similar stimuli evokes similar responses [2], [3]. Slightly abusing notation of *φ*, we use *φ*(**x**) and *φ*(`supp`(**x**)) interchangeably here. Smoothness is formally defined below.

#### Criterion 2 (Smoothness).

*The coding procedure between stimuli and their corresponding presynaptic encoded vectors is smooth if for the two representation vectors of two stimuli* **x** *and* **x**′, *we have* `supp`(*φ*(`supp`(**x**) ∩ `supp`(**x**′))) ⊆ `supp`(*φ*(**x**)) ∩ `supp`(*φ*(**x**′)).

The optimal information transmission is obtained when all information of a stimulus is transmitted with the minimum number of presynaptic neurons required to encode the stimulus in order to decode it exactly or approximately. The number of presynaptic neurons used to encode a stimulus must be at least equal to the information-theoretic bound. Otherwise, the stimulus cannot be encoded as it is supposed to be.

#### Criterion 3 (optimal information transmission).

*The optimal information transmission criterion is attained if t is minimized*.

The optimal energetic cost for a stimulus is achieved once the number of the presynaptic neurons used is minimum and only the presumed postsynaptic neurons representing that stimulus are activated.

#### Criterion 4 (Optimal energetic cost).

*Suppose that an energy unit is required for a neuron to spike. Then the optimal energetic cost for activating the presumed postsynaptic neurons in* `supp`(**x**) *is* min *t* + |`supp`(**x**)| *units*.

Finally, the low-dimensional representation and sparse coding are defined as follows.

#### Criterion 5 (Low-dimensional representation).

*A neural code is low-dimensional if for any stimulus, the number of postsynaptic neurons representing every stimulus is much smaller than the number of postsynaptic neurons. In other words, if that number is bounded by d then d* = *o*(*n*).

#### Criterion 6 (Sparse coding).

*The representation of a stimulus is sparse if the number of presynaptic neurons in response to the stimulus in the pursuit of activating exactly the presumed spiking postsynaptic neurons for the stimulus is much smaller than n, i*.*e*., *t* = *o*(*n*).

Normally, *t* would depend on *d* and *n*.

### B. Morphology of the brain

In this section, we show that to decode a stimulus with a few or no errors, the brain must have a certain morphology. Let the Boolean sum (or union) of *k* vectors **v**_1_, …, **v**_*k*_ of length *t* be 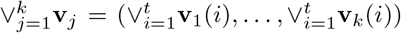, where ∨ is the bitwise OR operator. A vector **v** = (*v*_1_, …, *v*_*t*_) is not included in a vector **u** = (*u*_1_, …, *u*_*v*_) if there exists *i* ∈ [*t*] such that *v*_*i*_ = 1 and *u*_*i*_ = 0. In other words, `supp`(**v**) ⊈ `supp`(**u**). We define superimposed code as follows.

#### Definition 1.

*[47, Remark 1 and Definition 1] A t* × *n matrix M is a* superimposed (*d, n, r*)-code *of length t if the Boolean sum of any r column in M is not included in the Boolean sum of any other d columns. In other words, for any d* + *r distinct columns j*_1_, *j*_2_, …, *j*_*d*+*r*_, *one has* 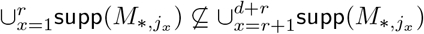.

We are now ready to state a fundamental result on the morphology of brain in the pursuit of the decoding objective as follows.

#### Theorem 1

(Morphology of the brain).*Consider the simplified model in the noiseless setting in Section II-C. Let M* = (*m*_*ij*_) *be a presynaptic-postsynaptic connectivity matrix, where m*_*ij*_ = 1 *means the presynaptic neuron i has a synaptic connection with the postsynaptic neuron j, and m*_*ij*_ = 0 *means otherwise. Criterion 1 holds if and only if M is a superimposed* (*d, n, r*)*-code*.

*Proof:* To prove this theorem, we have to precisely identify 𝒳, 𝒴, the encoding map *φ*, the decoding rule, and the built-in stimulus space. From the definition of *M*, we have 𝒮 _*i*_ = {*j* ∈ [*n*] | *m*_*ij*_ = 1}. We first prove the conditional statement, i.e., if Criterion 1 holds then *M* is a superimposed (*d, n, r*)-code, by contradiction.

*Conditional statement:* Assume that 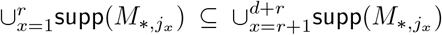. Consider the stimulus represented by *d* spiking postsynaptic neurons *j*_*r*+1_, …, *j*_*d*+*r*_. Because every postsynaptic neurons *j*\* ∈ {*j*_1_, …, *j*_*r*_} is not activated, there must exist a presynaptic neuron, say *i*\*, connecting with the postsynaptic neuron *j*\*, that expresses as an inhibitory neuron. In other words, *y*_*i*\*_ = 0 and *j*\* ∈ 𝒮_*i*\*_. On the other hand, because 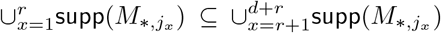, for every 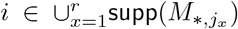, there exists 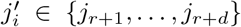 such that 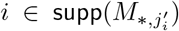. Therefore, there must exist some 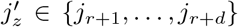 such that 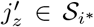. The postsynaptic neuron 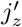 is then not activated because *y*_*i*\*_ = 0. This means the spiking postsynaptic neurons *j*_1_, …, *j*_*d*_ representing the stimulus cannot be activated as they are supposed to be. In other words, Criterion 1 does not hold.

When the stimulus is represented by less than *d*′ *< d* spiking postsynaptic neurons, the above arguments still hold because if 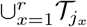 is not a subset of the union of 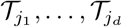 then it is also not a subset of the union of less than *d* sets in 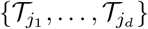.

*Converse:* We start by proving the converse by defining the encoding map. Given a stimulus represented by a status of the postsynaptic neurons, i.e., **x** ∈ 𝒳 ⊆ {0, 1}^*n*^, the received codeword **y** is given by

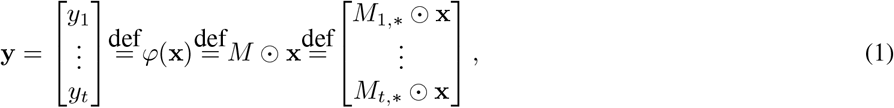

where *M*_*i*,*_ is the *i*th row of matrix *M*, ⊙ is a notation for the spiking rule; namely, *y*_*i*_ = *M*_*i*,*_ ⊙ **x** = 1 if 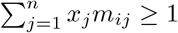, and *y*_*i*_ = *M*_*i*,*_ ⊙ **x** = 0 if 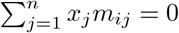 for *i* = 1, …, *t*. The encoding map is then defined as follows:

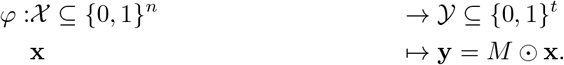

Consider the “naive” decoding algorithm to recover `supp`(**x**) from **y** = *M* ⊙ **x** as follows: keep removing items in 𝒮_*i*_ from [*n*] if *y*_*i*_ = 0 for *i* = 1, …, *t*. This algorithm is equivalent to the decoding rule of the decoder in the simplified model in the noiseless setting.

Let Dec(**y**, *M*) be the naive decoding procedure that takes **y** and *M* as the two inputs and produces a set of indices in [*n*]. When *M* is a superimposed (*d, n, r*)-code, it can be shown that Dec(*M* ⊙ **x**, *M*) = `supp`(**x**) ∪ ℒ if |`supp`(**x**)| ≤ *d*, where ℒ ⊆ [*n*] and |ℒ| ≤ *r* − 1 [48], [49]. Therefore, if *M* is a superimposed (*d, n, r*)-code then Criterion 1 holds. ▄

### C. Criterion analysis

In this section, we study causality and correlation between the six criteria described in Section III-A as illustrated in Figure 1.

#### 1) Lower bounds

Because a stimulus is encoded by *up to d* spiking postsynaptic neurons among a population of *n* postsynaptic neurons and each presynaptic neuron has two states, the minimum number of presynaptic neurons required to encode all stimuli must be at least 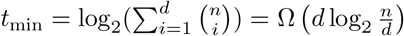 for exact decoding. This is the information-theoretic bound.

Suppose Criterion 1 holds. We now prove the lower bounds of *t* for exact decoding, i.e., *r* = 1, and approximate decoding, i.e., *r* ≥ 2.

#### Exact decoding

When *r* = 1, *M* becomes a *d*-disjunct matrix. It is well-known that 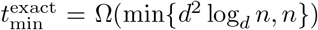 [47]. Therefore, it requires at least Ω(min{*d*^2^ log_*d*_ *n, n}*) presynaptic neurons to carry all information contained in the stimulus for exact decoding.

#### Approximate decoding

Cheraghchi [50] proves a lower bound of length of 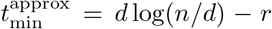 for any (*d, n, r*)-superimposed code. Therefore, it requires at least Ω(min{*d* log(*n/d*) − *r, n*}) presynaptic neurons to carry all information contained in the stimulus for approximate decoding.

It is noted that the number of rows of *M* can be achieved with *t* = *O*(min{*d*^2^ log_*d*_ *n, n*}) [51] for exact decoding and *t* = *O*(*d* log(*n/d*)) for approximate decoding with *r* = *d* [49], [52]. Since *d* has a direct impact on the magnitude of *t*, Criterion 5 correlates with Criterion 3.

When Criterion 3 is obtained, Criterion 4 is automatically obtained for exact decoding case. Moreover, when *r* = *d*, the number of presynaptic neurons obtains the information-theoretic by allowing up to *d* − 1 additional spiking postsynaptic neurons along with the postsynaptic neurons representing a stimulus. In addition, for some stimuli, exact decoding can be obtained by chance. This trade-off is fairly acceptable. We summarize the discussion above by the following lemma.

##### Lemma 1.

*Consider the simplified model in the noiseless setting in Section II-C. Suppose Criterion 1 holds. It requires at least* 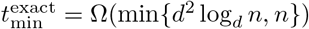 *(respectively*, 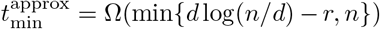*) postsynaptic neurons spiking to encode an input stimulus for exact decoding (respectively, approximate decoding). Moreover, if* 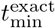 *in exact decoding case is obtained then the optimal energetic cost, i*.*e*., *Criterion 4, is also obtained*.

#### 2) Smoothness

This section is devoted to prove that smoothness in Criterion 2 is caused by Criterion 1.

##### Lemma 2.

*Consider the simplified model in the noiseless setting in Section II-C. If Criterion 1 holds then Criterion 2 also holds*.

*Proof:* From the definition of **y** in (1), we have

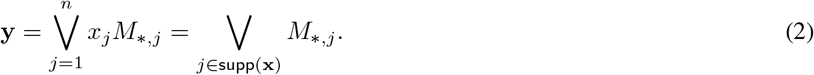

Consider two stimuli **x** and **x**′. Set ℛ = `supp`(**x**) ∩ `supp`(**x**′). By using (2), the two encoded codewords of the two stimuli are:

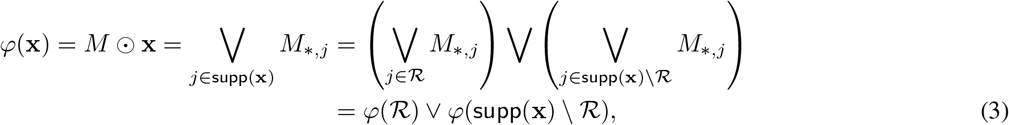

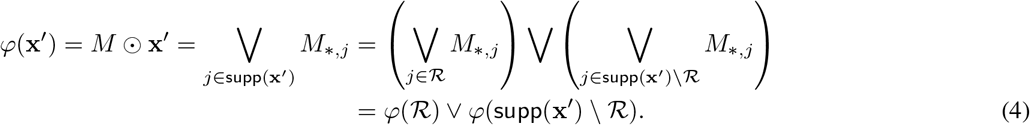

Because of the bitwise OR operator, we get `supp`(*φ*(**x**)) = `supp`(*φ*(ℛ)) ∪ `supp`(*φ*(`supp`(**x**) \ ℛ)) and `supp`(*φ*(**x**′)) = `supp`(*φ*(ℛ)) ∪`supp`(*φ*(`supp`(**x**′)\ ℛ)). This implies `supp`(*φ*(ℛ)) = `supp`(*φ*(`supp`(**x**) ∩`supp`(**x**′))) ⊆ `supp`(*φ*(**x**)) ∩`supp`(*φ*(**x**′)).

▄

#### 3) Low-dimension representation and sparse coding

Suppose Criteria 1 and 3 hold. If Criterion 5 holds for 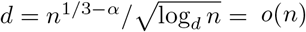 with *α <* 1*/*3, we must have *t* = *O*(max{min{*d*^2^ log_*d*_ *n, n*}, min{*d* log(*n/d*) −*r, n*}}) = *O*(*d*^2^ log_*d*_ *n*) = *o*(*n*). Therefore, Criterion 6 holds. If Criterion 6 holds, i.e., *t* = *o*(*n*), we must have

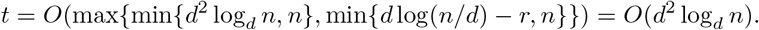

It is obvious that *d* = *o*(*n*). We summarize these result below.

##### Lemma 3.

*Consider the simplified model in the noiseless setting in Section II-C. Suppose Criteria 1 and 3 hold. If Criterion 5 holds for* 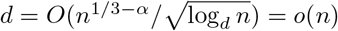 *with α <* 1*/*3 *then Criterion 6 also holds. On the other hand, if Criterion 6 holds then Criterion 5 holds*.

## IV. Testing method

To attain the decoding rule in biological neural networks, we assume that every presynaptic neuron is hybrid as in Section II-A1. This might be too ideal to be true in practice, though we suggest that synaptic integration and synergy could resolve this assumption in practice. As discussed in Section I, if the proposed model is satisfied, by varying *d* and setting *t* as optimal information transmission accordingly such that *t* = *o*(*n*), the probability of matrix matching is equal to the probability of hypothesis matching. However, if the simplified model in the noiseless setting is not satisfied, one can only test the probability of matrix matching.

### A. Matrix instantiation

As in Section III-C1, for exact (respectively, approximate) decoding, i.e., *r* = 1 (respectively, *r* ≥ 2), *M* becomes a *d*-disjunct matrix (respectively, a superimposed (*d, n, r*)-code). Disjunct matrices are defined as follows.

#### Definition 2.

*Given integers d and n, with* 1 ≤ *d* ≤ *n, we say that a t* × *n Boolean matrix M is a d-disjunct matrix if any submatrix of M obtained by choosing d*+1 *out of n arbitrary columns of M contains every distinct rows of the* (*d*+1) × (*d*+1) *identity matrix*.

Therefore, we have a simple procedure to check whether a *t* × *n* matrix *M* is a *d*-disjunct.

#### Remark 1.

*For every d* + 1 *columns in M, if they do not contain every distinct rows of the* (*d* + 1) × (*d* + 1) *identity matrix then M is not d-disjunct. Otherwise, M is a d-disjunct matrix*.

Du and Hwang [53] prove an (achievable) upper bound on *t* for any *t* × *n d*-disjunct matrix as below.

#### Theorem 2.

*[53] Given* 1 ≤ *d < n. There exists a nonrandom t* × *n d-disjunct matrix with t <* 4*d*^2^ log *n* = *O*(*d*^2^ log *n*).

There are many types of superimposed codes. Here we are interested in a class of them called *selectors* [52] in which a (*d* + *r, r* + 1, *n*)-selector is a superimposed (*d, n, r*)-code.

#### Definition 3.

*Given integers d, r, and n, with* 1 ≤ *d, r* ≤ *n, we say that a t* × *n Boolean matrix M is a* (*d* +*r, r* +1, *n*)*-selector if any submatrix of M obtained by choosing d* + *r out of n arbitrary columns of M contains at least r* + 1 *distinct rows of the identity matrix I*_*d*+*r*_.

The main reason why we are interested in selectors because they can be extended to construct a (*d* + *r*)-disjunct matrix by adding *d* additional rows of the identity matrix *I*_*d*+*r*_.

Cheraghchi [50] proves a lower bound of length of *d* log(*n/d*) − *r* for any superimposed (*d, n, r*)-code. When *r* = Θ(*d*), any construction for (*d* + Θ(*d*), Θ(*d*), *n*)-selectors achieving the number of length *O*(*d* log(*n/d*)) is information-theoretically optimal. Bonis et al. [52] show that this bound can be achieved by setting *r* = *d*.

#### Theorem 3.

*[52] Given* 1 ≤ *d < n. There exists a t* × *n* (2*d, d* + 1, *n*)*-selector with* 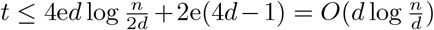.

Based on Definition 3, we have a simple procedure to check whether a *t* × *n* matrix *M* is a (2*d, d* + 1, *n*)-selector below.

#### Remark 2.

*For every* 2*d columns in M, if they do not contain at least d* + 1 *distinct rows of the identity matrix I*_2*d*_ *then* 𝒮 *is not a* (2*d, d* + 1, *n*)*-selector. Otherwise, M is a* (2*d, d* + 1, *n*)*-selector*.

### B. Testing of population proportion

We would like to calculate a proportion of successful trials in a population of trials. However, since the population is too large to obtain its census, we can only observe a fraction of the population including some successful trials. The question is to find the confidence interval with a given confidence level for a proportion is similar to that for the population mean from the observable population of trials.

If `X` is a binomial random variable, then `X` ∼ 𝔹(*m, p*), where *m* is the number of trials obtained from the population and *p* is the probability of a successful trial. The estimated proportion is defined as

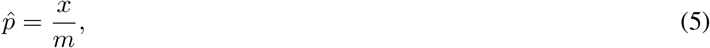

where *x* is the count of successes in the sample of *m* trials.

Then a population proportion can be estimated through the usage of a confidence interval known as a one-sample proportion in the *z*-interval as follows:

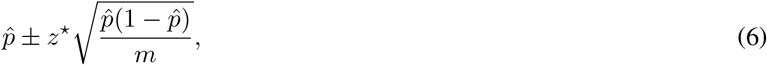

where 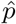 is defined in (5), *m* is the number of trials, and *z*^⋆^ is the upper (1 − *c*)*/*2 critical value of the standard normal distribution for a level of confidence *c* (*z*^⋆^ = 1.96 if *c* = 0.95). The parameter 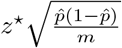 is called the error bound for the proportion 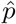.

### C. Probability of matrix matching

Our goal is to verify whether the presynaptic-postsynaptic connectivity matrix *M* is a *d*-disjunct matrix or a (2*d, d* + 1, *n*)- selector for 2 ≤ *d*. The complexity of testing whether *M* is a *d*-disjunct matrix (respectively, a (2*d, d* + 1, *n*)-selector) is 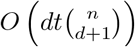 (respectively, 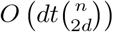) because we have to scan every *d* + 1 (respectively, 2*d*) columns out of *n* columns in *M*. Therefore, it is nearly impossible to complete such task with the current resources. We propose an alternative way to test whether *M* is a *d*-disjunct matrix or a (2*d, d* + 1, *n*)-selector based on population proportion testing. The procedure is as follows.

To test whether *M* is a *d*-disjunct matrix, we first select *m* = 100, 000 distinct trials in which each trial is a selection of *d* + 1 columns out of *n* columns in *M* to test whether they contain the identity matrix *I*_*d*+1_. A successful trial is called if those *d* + 1 columns contain an identify matrix *I*_*d*+1_. The probability of *M* being *d*-disjunct belongs to the range defined in (6). This is also the probability of matrix matching.

Since the number of trials is *m* = 100, 000 and the confidence level is 95%, the error bound for our proportion is up to

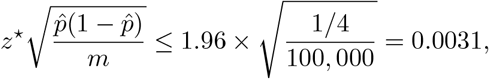

which is negligible. Therefore, *we can assign* 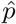 *as the probability of matrix matching*.

Similarly, the procedure to test whether *M* is a (2*d, d* + 1, *n*)-selector by taking each trial as a selection of 2*d* columns out of *n* columns in *M* to test whether they contain at least *d* + 1 distinct rows of the identity matrix *I*_2*d*_. A successful trial occurs when the query is true.

In summary, we defined the probability of matrix matching as follows.

#### Definition 4.

*Let M be a t* × *n input matrix. To verify whether M is a d-disjunct, a trial is a selection of d*+1 *columns out of n columns in M to test whether they contain the identity matrix I*_*d*+1_. *Meanwhile, to verify whether M is a* (2*d, d* +1, *n*)*-selector, a trial is a selection of* 2*d columns out of n columns in M to test whether they contain at least d* + 1 *distinct rows of the identity matrix I*_2*d*_. *A trial is successful if the query is true*.

*Let m* = 100, 000 *and x be the numbers of total trials and successful trials, respectively, for testing whether M is a d-disjunct or* (2*d, d* + 1, *n*)*-selector. Then the probability of matrix matching is* 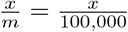.

## V. Experimental results

### A. Functional connectivity network

#### 1) Description

The original dataset is obtained from [54] as a 638 × 638 square network matrix *T* = (*t*_*ij*_) in which the rows and columns in the matrix represent network nodes and matrix entries represent node connectivity strength. The authors construct a functional connectivity network matrix from correlations between regional fMRI times series, measured at the same 638 nodal locations as in the coactivation analysis, from 27 healthy volunteers scanned in the resting state. Each entry is calculated by using the Jaccard index as the metric of functional coactivation strength, defined for each pair of regions as the number of contrasts activating both regions *X* and *Y* divided by the union of contrasts activating region *X* and activating region *Y*. Entry *t*_*ij*_ is larger than 0 means there is a connection between node *i* and node *j*, and entry *t*_*ij*_ is equal to 0 means otherwise.

#### 2) Preprocessing

We convert the matrix *T* into a binary matrix *G* = (*g*_*ij*_) in which *g*_*ij*_ = 1 if *t*_*ij*_ *>* 0 and *g*_*ij*_ = 0 otherwise.

Consider 638 nodes as 638 neurons. We then create a *t* × *n* binary presynaptic-postsynaptic connectivity matrix *M* = (*m*_*ij*_) as follows, where *n* is the number of postsynaptic neurons and *t* is the number of presynaptic neurons. Let 𝒮 ⊆ [638] = {1, …, 638} with |𝒮| = *t* be a set of presynaptic neurons and 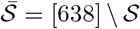 be the set of postsynaptic neurons corresponding to the set of presynaptic neurons 𝒮. Matrix *M* is obtained by removing every column *G* _*\*,j*_ and every row *G*_*i,\**_ in *G*, for *j* ∈ *S* and 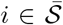. It is easy to confirm that the new matrix *M* = (*m*_*ij*_) obtained from the removing procedure has size of *t* × *n*. Entry *m*_*ij*_ = 1 means there is a connection between the presynaptic neuron *i* and the postsynaptic neuron *j*, and *m*_*ij*_ = 0 means otherwise.

#### 3) Experiments

Because of the preprocessing in the previous subsection, each presynaptic neuron can be set to be hybrid. Therefore, the simplified model in the noiseless setting is satisfied. As discussed in the beginning of this section, the probability of matrix matching is then equal to the probability of hypothesis matching.

Because of the lower bound on (2*d, d* + 1, *n*)-selectors, we set the lower bound for *t* as 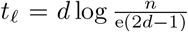. When testing whether *M* is a *d*-disjunct, the upper bound for *t* is *d*^2^ log *n*. We also set the upper bound for *t* as 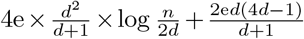 as in Theorem 3 when testing whether *M* is a (2*d, d* + 1, *n*)-selector.

We run experiments for *d* = {4, 8, 15} to get the probability of hypothesis matching, which is also the probability of matrix matching defined in Definition 4. The result is plotted in Figure 4. The red and green lines represent testing whether *M* is a *d*-disjunct matrix or (2*d, d* + 1, *n*)-selector, respectively.

**Fig. 4.**
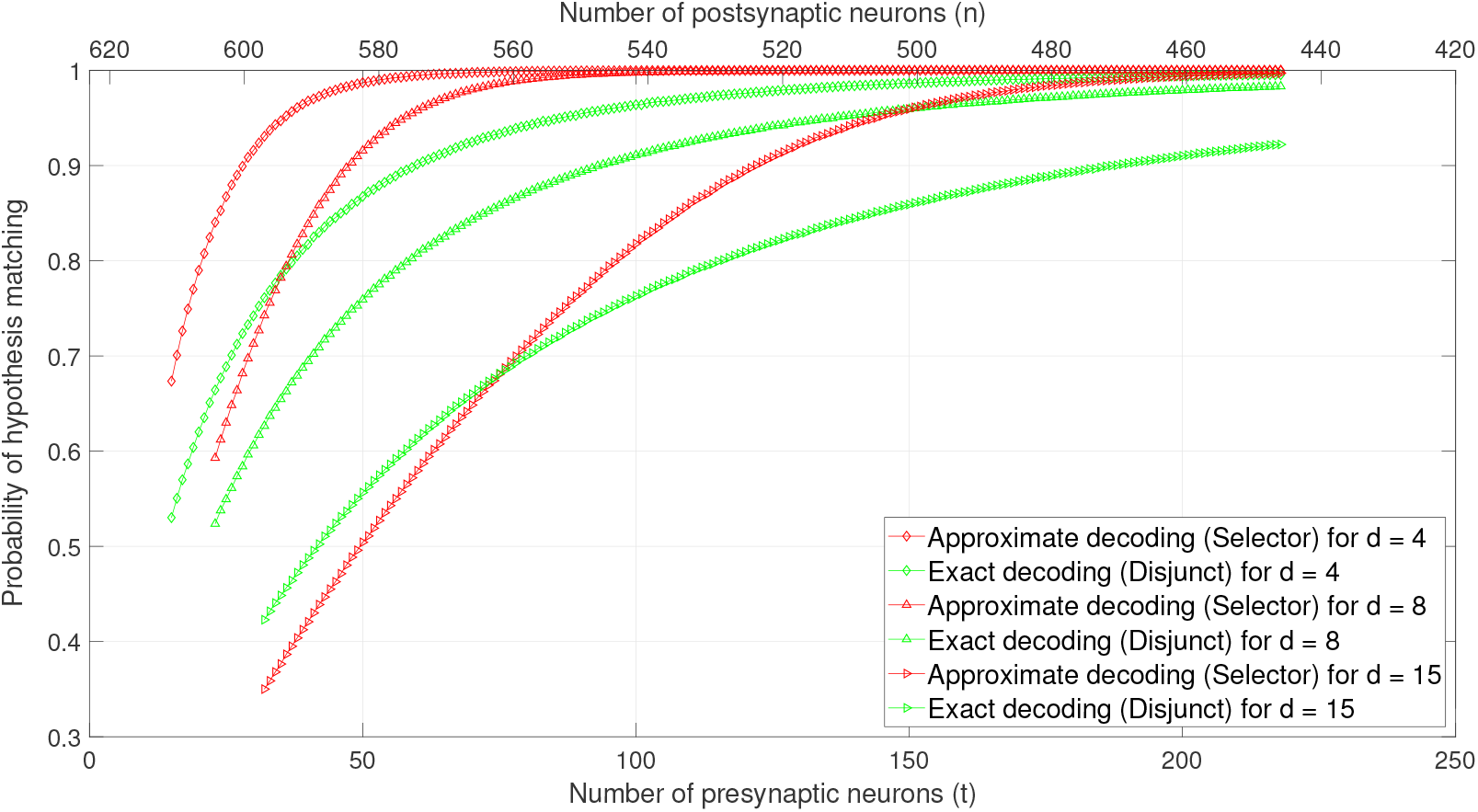
Probability of hypothesis matching by varying *d* (the low-dimensional representation), the number of presynaptic neurons (sparse coding), and the population of postsynaptic neurons.

As we can see, once *d* is small, the probability of hypothesis matching converges quickly to 1 when the ratio between the number of presynaptic neurons and the number of postsynaptic neurons is smaller than 138*/*500 = 0.276. However, once *d* grows, the probability that *M* is a *d*-disjunct does not converge to 1. This phenomenon can be explained by the requirement of a large number of presynaptic neurons compared to the number of postsynaptic neurons. Interestingly, the probability that *M* is a (2*d, d* + 1, *n*)-selector still converges to 1. Moreover, once *M* is a selector, the optimal latency, smoothness, and low-dimensional and high-dimensional representations, and the optimal energetic cost are obtained while optimal information transmission almost holds. This highly suggests that **selectors may be the right topology for the presynaptic-postsynaptic connectivity in brain**.

### B. Layer 2 of the medial entorhinal cortex

The dataset is obtained from a 25-day-old rat [55] in layer 2 of the medial entorhinal cortex. It is represented as an 18 × 180 presynaptic-posynaptic connectivity matrix *M* = (*m*_*ij*_) in which the rows and columns represent presynaptic and postsynaptic neurons, respectively. Entry *m*_*ij*_ = 1 means there is a connection between the *i*th presynaptic neuron and the *j*th postsynaptic neuron. There are 15 excitatory and 3 inhibitory presynaptic neurons in the presynaptic site while the population of postsynatic neurons includes 102 excitatory and 78 inhibitory neurons.

Since the classification of the presynaptic neurons is determined, we are unable to verify whether the simplified model in the noiseless setting is satisfied. Therefore, we only test the probability of matrix matching for matrix *M*. We run experiments for *d* = 2 to *d* = 15 to get the probability of matrix matching as defined in Definition 4.

As plotted in Figure 5, the probability that *M* is either a *d*-disjunct matrix or a (2*d, d* + 1, *n*)-selector decreases as *d* increases. The main reason is because the number presynaptic neurons is too small and the number of postsynaptic neurons is also relatively small. However, even with this small dataset, a positive sign is that *M* still is almost 2-disjunct or (4, 3, 180)- selector when *d* = 2. This indicates that our suggestion that synergy could make our proposed model work in reality may be true.

**Fig. 5.**
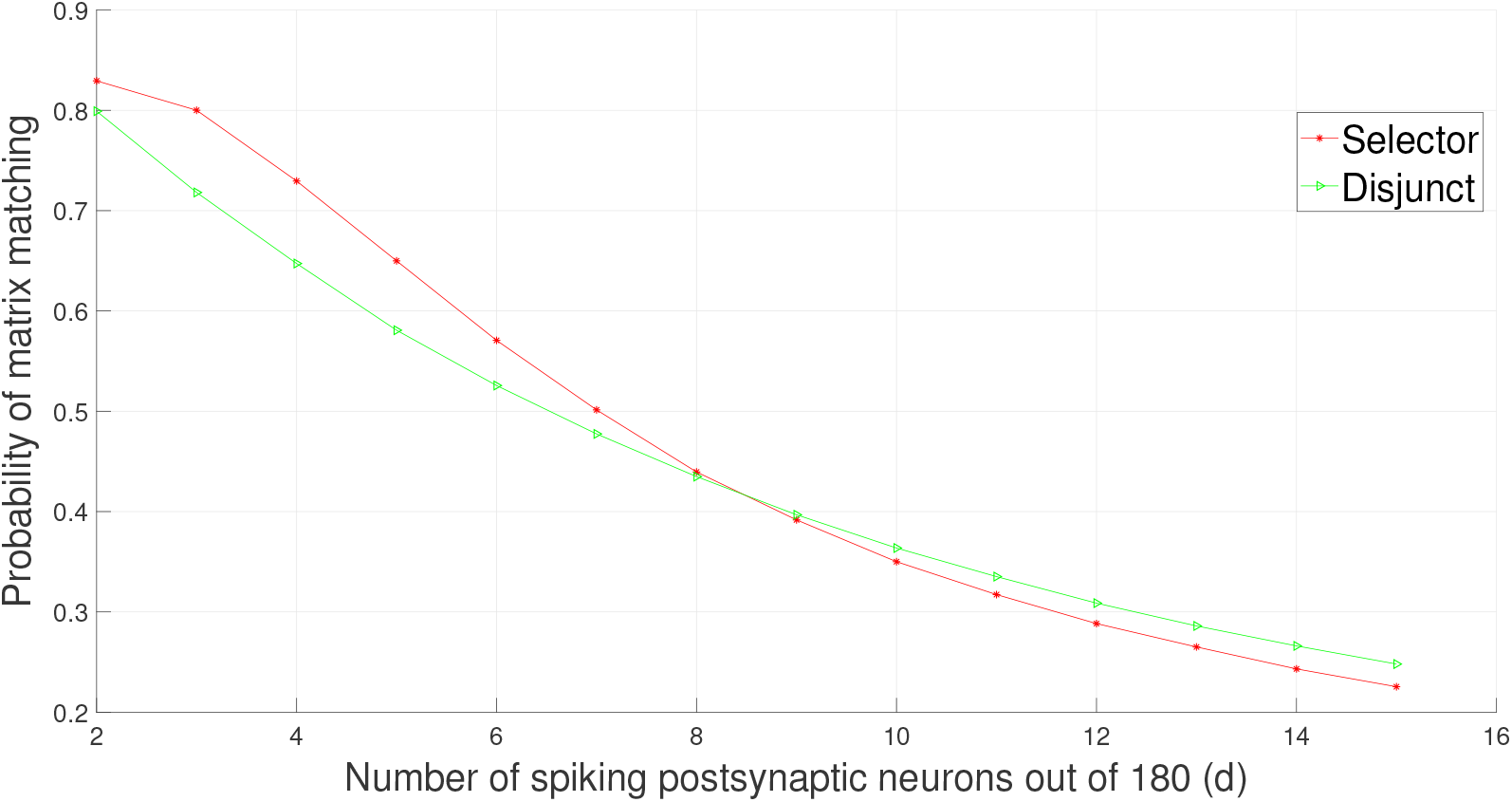
Probability of matrix matching for the 18 × 180 presynaptic-posynaptic connectivity matrix (*t* = 18, *n* = 180).

